# Contrasting regional architectures of schizophrenia and other complex diseases using fast variance components analysis

**DOI:** 10.1101/016527

**Authors:** Po-Ru Loh, Gaurav Bhatia, Alexander Gusev, Hilary K Finucane, Brendan K Bulik-Sullivan, Samuela J Pollack, Schizophrenia Working Group of the Psychiatric Genomics Consortiumy, Teresa R de Candia, Sang Hong Lee, Naomi R Wray, Kenneth S Kendler, Michael C O’Donovan, Benjamin M Neale, Nick Patterson, Alkes L Price

**Affiliations:** Department of Epidemiology, Harvard T.H. Chan School of Public Health, Boston, Massachusetts, USA; Program in Medical and Population Genetics, Broad Institute of Harvard and MIT, Cambridge, Massachusetts, USA; Department of Mathematics, Massachusetts Institute of Technology, Cambridge, Massachusetts, USA; Analytic and Translational Genetics Unit, Massachusetts General Hospital, Boston, Massachusetts, USA; Department of Biostatistics, Harvard T.H. Chan School of Public Health, Boston, Massachusetts, USA; Department of Psychology and Neuroscience, University of Colorado Boulder, Boulder, Colorado, United States; The Queensland Brain Institute, University of Queensland, Brisbane, Queensland, Australia; Department of Psychiatry and Human Genetics, Virginia Institute of Psychiatric and Behavioral Genetics, Virginia Commonwealth University, Richmond, Virginia, USA; MRC Centre for Neuropsychiatric Genetics and Genomics, Institute of Psychological Medicine and Clinical Neurosciences, Cardiff University, Cardiff, UK

## Abstract

Heritability analyses of GWAS cohorts have yielded important insights into complex disease architecture, and increasing sample sizes hold the promise of further discoveries. Here, we analyze the genetic architecture of schizophrenia in 49,806 samples from the PGC, and nine complex diseases in 54,734 samples from the GERA cohort. For schizophrenia, we infer an overwhelmingly polygenic disease architecture in which ≥71% of 1Mb genomic regions harbor at least one variant influencing schizophrenia risk. We also observe significant enrichment of heritability in GC-rich regions and in higher-frequency SNPs for both schizophrenia and GERA diseases. In bivariate analyses, we observe significant genetic correlations (ranging from 0.18 to 0.85) among several pairs of GERA diseases; genetic correlations were on average 1.3x stronger than correlations of overall disease liabilities. To accomplish these analyses, we developed a fast algorithm for multi-component, multi-trait variance components analysis that overcomes prior computational barriers that made such analyses intractable at this scale.

---

Over the past five years, variance components analysis has had considerable impact on research in human complex trait genetics, yielding rich insights into the heritable phenotypic variation explained by SNPs [1–3], its distribution across chromosomes, allele frequencies, and functional annotations [4–6], and its correlation across traits [7,8]. These analyses have complemented genome-wide association studies (GWAS): while GWAS have identified individual loci explaining significant portions of trait heritability, variance components methods have aggregated signal across large SNP sets, revealing information about polygenic SNP effects invisible to association studies. The utility of both approaches has been particularly clear in studies of schizophrenia, for which early GWAS achieved few genome-wide significant findings, yet variance components analysis indicated a large fraction of heritable variance spread across common SNPs in numerous loci, over 100 of which have now been discovered in large-scale GWAS [5,9–12].

Despite these advances, much remains unknown about the genetic architecture of schizophrenia and other complex diseases. For schizophrenia, known GWAS loci are collectively estimated to explain only 3% of variation in disease liability [12]; of the remaining variation, a sizable fraction has been shown to be hidden among thousands of common SNPs [5,11], but the distribution of these SNPs across the genome and across the allele frequency spectrum has remained uncertain. Even for traits such as lipid levels and type 2 diabetes for which loci of somewhat larger effect have been identified, the spatial and allelic distribution of variants responsible for the bulk of known SNP-heritability has remained a mystery [13,14]. Variance components methods have the potential to shed light on these questions using the increased statistical resolution offered by tens or hundreds of thousands of samples [15,16]. However, while study sizes have increased beyond 50,000 samples, existing variance components methods [2] are becoming computationally intractable at such scales. Computational limitations have thus forced previous studies to split and then meta-analyze data sets [6], a procedure that results in loss of precision for variance components analysis, which relies on pairwise relationships for inference (in contrast to meta-analysis in association studies) [15,16].

Here, we introduce a much faster variance components method, BOLT-REML, and apply it to analyze roughly 50,000 samples in each of two very large data sets—from the Psychiatric Genomics Consortium (PGC2) [12] and the Genetic Epidemiology Research on Aging cohort (GERA; see URLs)—obtaining several new insights into the genetic architectures of schizophrenia and nine other complex diseases. We harnessed the computational efficiency and versatility of BOLT-REML variance components analysis to estimate components of heritability, infer levels of polygenicity, partition SNP-heritability across the common allele frequency spectrum, and estimate genetic correlations among GERA diseases. We corroborated our results using an efficient implementation of PCGC regression [17] when computationally feasible to do so.

## Results

### Overview of Methods

The BOLT-REML algorithm employs the conjugate gradient-based iterative framework for fast mixed model computations [18,19] that we previously harnessed for mixed model association analysis using a single variance component [20]. In contrast to that work, BOLT-REML robustly estimates variance parameters for models involving multiple variance components and multiple traits [21,22]. BOLT-REML uses a Monte Carlo average information restricted maximum likelihood (AI REML) algorithm [23], which is an approximate Newton-type optimization of the restricted log likelihood [24] with respect to the variance parameters being estimated. (In contrast, our previous work [20] used a rudimentary quasi-Newton approach that sufficed only for univariate optimization.) In each iteration, BOLT-REML rapidly approximates the gradient of the log likelihood using pseudorandom Monte Carlo sampling [25] and approximates the Hessian of the log likelihood using the average information matrix [26]. Full details, including simulations verifying the accuracy of BOLT-REML heritability parameter estimates and standard errors (which are nearly identical to standard REML), are provided in Online Methods and the Supplementary Note. We have released open-source software implementing the method (see URLs).

### Computational efficiency of BOLT-REML variance components analysis

We assessed the computational performance of BOLT-REML, comparing it to the GCTA software [2] (see URLs) for REML variance components analyses of GERA disease phenotypes on subsets of the GERA cohort of increasing size. We observed that across three types of analyses, BOLT-REML achieved order-of-magnitude reductions in running time and memory use compared to GCTA, with relative improvements increasing with sample size (Figure 1). The running times we observed for BOLT-REML scale roughly as *≈M N*^1.5^, consistent with previously reported empirical results for BOLT-LMM association analysis [20], whereas standard REML analysis requires *O*(*M N*^2^ + *N*^3^) running time (Figure 1a and Supplementary Table 1). BOLT-REML also only requires *≈M N/*4 bytes of memory (nearly independent of the number of variance components used), in contrast to standard REML analysis, which requires *O*(*N*^2^) memory per variance component (Figure 1b and Supplementary Table 1). Consequently, GCTA could only analyze at most half of our available samples; indeed, computational constraints have forced previous studies to split large cohorts into multiple subgroups for analysis [6], increasing standard errors and reducing statistical power. In contrast, BOLT-REML enabled us to perform a full suite of heritability analyses of *N* =50,000 samples with tight error bounds [15,16].

**Figure 1.**
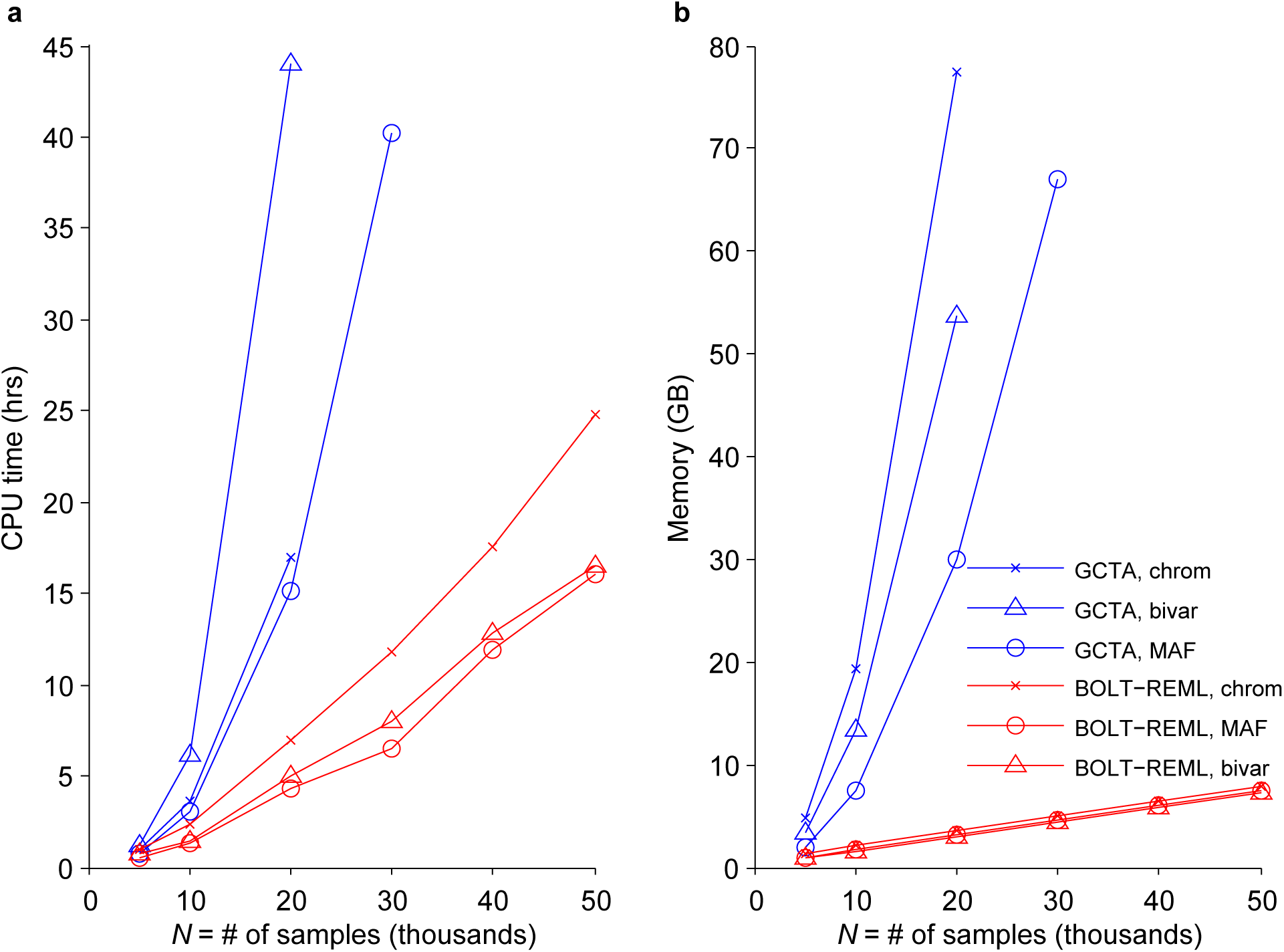
Computational performance of BOLT-REML and GCTA heritability analysis algorithms. Benchmarks of BOLT-REML and GCTA in three heritability analysis scenarios: partitioning across 22 chromosomes, partitioning across six MAF bins, and bivariate analysis. Run times (**a**) and memory (**b**) are plotted for runs on subsets of the GERA cohort with fixed SNP count *M* =597,736 and increasing sample size (*N*) using dyslipidemia as the phenotype in the univariate analyses and hypertension as the second phenotype in the bivariate analysis. Reported run times are medians of five identical runs using one core of a 2.27 GHz Intel Xeon L5640 processor. Reported run times for GCTA are total times required for computing the GRM and performing REML analysis; time breakdowns and numeric data are provided in Supplementary Table 1. Data points not plotted for GCTA indicate scenarios in which GCTA required more memory than the 96GB available. Software versions: BOLT-REML, v2.1; GCTA, v1.24.

### Estimates of SNP-heritability for schizophrenia and GERA diseases

We analyzed 22,177 schizophrenia cases and 27,629 controls with well-imputed genotypes at 472,178 markers of minor allele frequency (MAF) *≥*2% in the PGC2 data [12] (Supplementary Table 2) as well as nine complex diseases in 54,734 randomly ascertained samples typed at 597,736 SNPs in the GERA cohort (see Online Methods; QC procedures included filtering both data sets to unrelated samples of European ancestry and LD-pruning markers to *r*^2^*≤*0.9). To remove possible effects of population stratification, all analyses included 10 principal component covariates and PGC2 analyses further included 29 study indicators (see Online Methods). We computed liability-scale SNP-heritability estimates (*h*_g_^2^, ref. [1]) for schizophrenia in the PGC2 data set and all 22 disease phenotypes in the GERA data set assuming a liability threshold model; we assumed schizophrenia population risk of 1% (ref. [5,11,12]), and we assumed population risks of GERA diseases matched case fractions in the GERA cohort. For the GERA diseases, we estimated *h*_g_^2^ by applying BOLT-REML directly to observed case/control status—obtaining raw observed-scale heritability parameter estimates *h*_g–cc_^2^ —and then converting *h*_g–cc_^2^ to liability-scale *h*_g_^2^ using the linear transformation of ref. [3] (Table 1 and Supplementary Table 3). Given the very low values of *h*_g–cc_^2^ for many GERA diseases, we restricted further GERA analyses to the nine individual diseases with highest *h*_g–cc_^2^ (Table 1). For schizophrenia, we estimated *h*_g_^2^ by developing and applying a computationally efficient implementation of PCGC regression [17] (see URLs and Online Methods) in light of the known downward bias of large-sample REML *h*_g_^2^ estimates for ascertained casecontrol traits [17,27]. Indeed, upon performing REML analyses on full data sets as well as on subsamples of each data set with 2x–10x fewer samples, we observed significant downward bias of schizophrenia *h*_g_^2^ estimates with increasing sample size, whereas we observed no such trend for data from GERA, which is a cohort study not subject to case-control ascertainment (Supplementary Table 4). REML *h*_g_^2^ estimates on 10x-downsampled (*N ≈*5,000) PGC2 data corroborated the PCGC regression estimate (Supplementary Table 4), but we believe that PCGC regression is the most appropriate method for estimating genome-wide *h*_g_^2^ in ascertained case-control data.

**Table 1.**
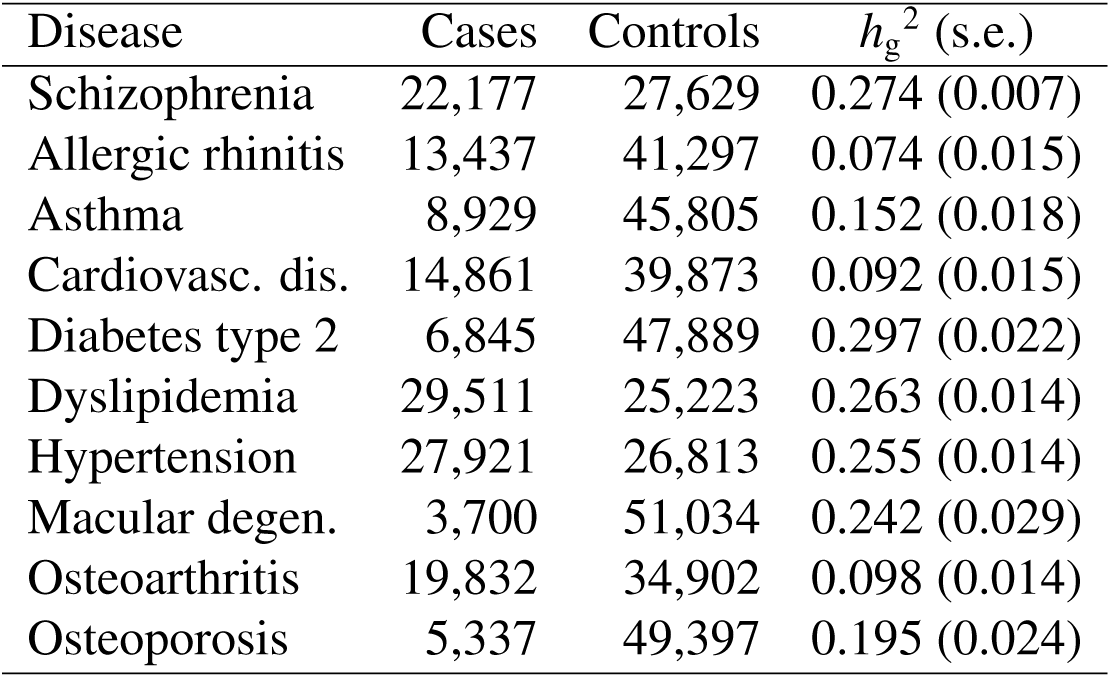
Estimated proportions of variance in disease liability explained by SNPs. Schizophrenia cases and controls are from the PGC2 data set [12]; the *h*_g_^2^ estimate assumes a population risk of 1% and was computed using PCGC regression to avoid REML bias induced by over-ascertainment of cases [17,27]. Cases and controls for the other 9 diseases are from the GERA data set; *h*_g_^2^ estimates assume random sample ascertainment and were computed using BOLT-REML.

| Disease | Cases | Controls | $h_g^2$ (s.e.) |
| --- | --- | --- | --- |
| Schizophrenia | 22,177 | 27,629 | 0.274 (0.007) |
| Allergic rhinitis | 13,437 | 41,297 | 0.074 (0.015) |
| Asthma | 8,929 | 45,805 | 0.152 (0.018) |
| Cardiovasc. dis. | 14,861 | 39,873 | 0.092 (0.015) |
| Diabetes type 2 | 6,845 | 47,889 | 0.297 (0.022) |
| Dyslipidemia | 29,511 | 25,223 | 0.263 (0.014) |
| Hypertension | 27,921 | 26,813 | 0.255 (0.014) |
| Macular degen. | 3,700 | 51,034 | 0.242 (0.029) |
| Osteoarthritis | 19,832 | 34,902 | 0.098 (0.014) |
| Osteoporosis | 5,337 | 49,397 | 0.195 (0.024) |

These analyses help explain a previously mysterious observation of decreasing estimated schizophrenia *h*_g_^2^ with increasing aggregation of cohorts [5]. This phenomenon was attributed to phenotypic heterogeneity [5,11], as suggested by estimates of between-cohort genetic correlation *<*1 (ref. [5]). Our analyses implicate ascertainment-induced downward bias of estimated *h*_g_^2^ (worsening with sample size) as an additional explanation of this effect (Supplementary Tables 4 and 5). In theory, the extent of ascertainment-induced bias could be used to infer the extent of case over-ascertainment and hence infer population risk, but we found in simulations that larger sample sizes would be required (Supplementary Table 6). Finally, we note that while our reported schizophrenia *h*_g_^2^ assumes a population risk of 1% (ref. [5,11,12]), this assumption does not affect estimates of the relative partitioning of SNP-heritability across SNP subsets; in the partitioning analyses that follow, *h*_g_^2^ serves only as a scale factor (Online Methods). Similarly, while our use of an LD-pruned marker set to alleviate LD bias [28–30] (Online Methods) results in a higher *h*_g_^2^ estimate than using unpruned markers (Supplementary Table 5), this choice does not otherwise affect the analyses that follow.

### Contrasting polygenicity of schizophrenia and GERA diseases

We next turned to a detailed investigation of the polygenicity of schizophrenia and the GERA diseases. Specifically, we estimated SNP-heritability explained by each 1Mb region of the genome, *h*_g,1Mb_^2^ (defined in Online Methods; Fig. 2a); we confirmed in simulations that 1Mb regions are sufficiently wide to ensure negligible leakage of heritability across region boundaries due to linkage disequilibrium or incomplete tagging of variants (Supplementary Tables 7 and 8). We restricted our primary analyses of GERA diseases to dyslipidemia and hypertension, the diseases with the highest observed-scale SNP-heritability *h*_g–cc_^2^ (Supplementary Table 3), because we had insufficient statistical power to make inferences for diseases with lower *h*_g–cc_^2^ (Supplementary Fig. 1). As expected, SNP-heritability estimates for individual 1Mb regions were individually noisy (mean estimated *h*_g,1Mb_^2^ / mean s.e. *h*_g,1Mb_^2^ for schizophrenia and 0.51 for dyslipidemia and hypertension), although we did see substantial SNP-heritability in some 1Mb regions (particularly for dyslipidemia, which has relatively large-effect SNPs [13]; in contrast, no 1Mb region was estimated to explain more than 0.1% of schizophrenia liability). We therefore sought to draw inferences from the bulk distribution of per-megabase SNP-heritability estimates (Supplementary Fig. 2). (We note that a limitation of BOLT-REML is that it is does not compute likelihood ratio test statistics for testing whether individual variance components contribute nonzero variance; see Supplementary Note.)

**Figure 2.**
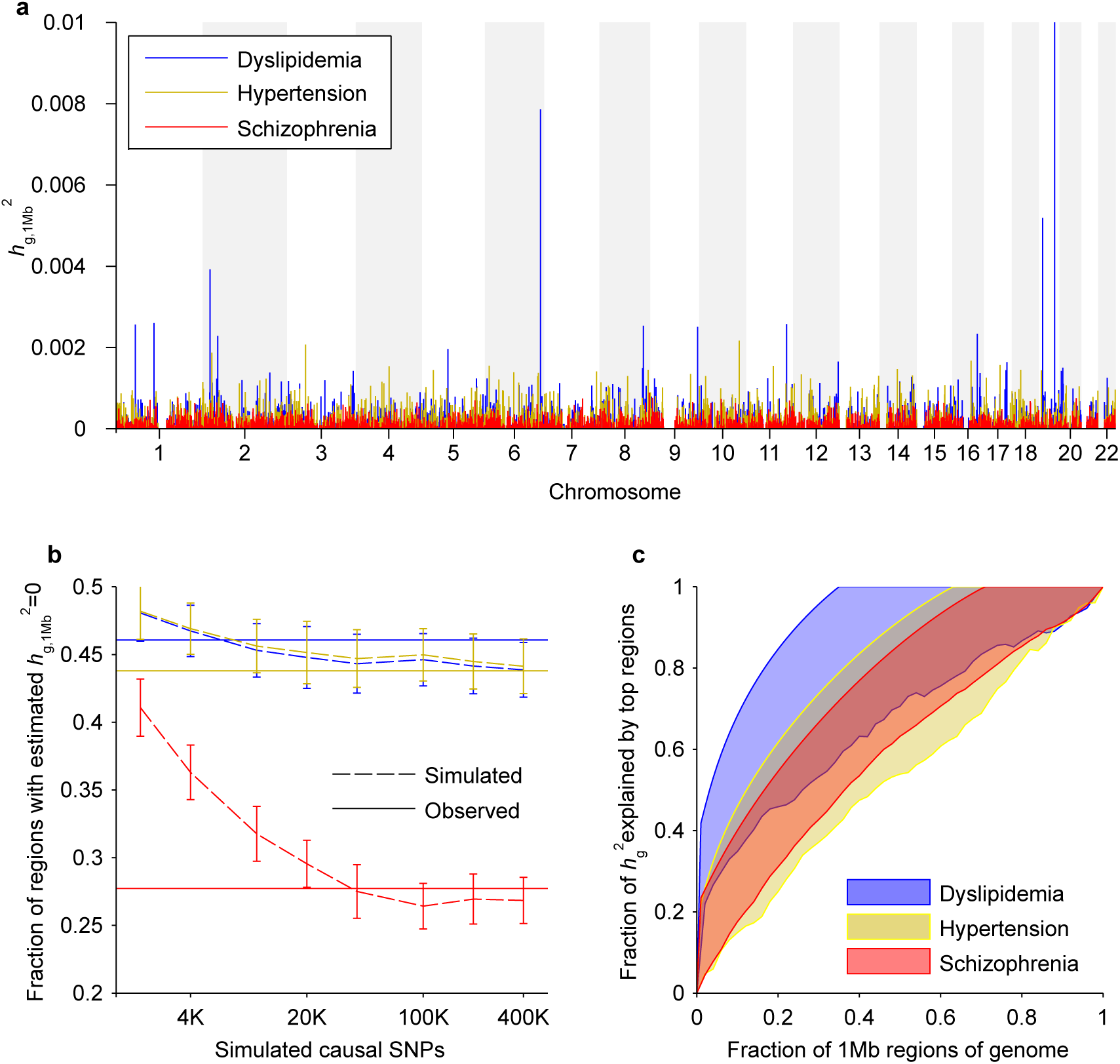
Extreme polygenicity of schizophrenia compared to other complex diseases. (**a**) Manhattan-style plots of estimated SNP-heritability per 1Mb region of the genome, *h*_g,1Mb_^2^, for dyslipidemia, hypertension, and schizophrenia. The *APOE* region of chromosome 19 is an outlier with an *h*_g,1Mb_^2^ estimate of 0.022. (**b**) Fractions of 1Mb regions with estimated *h*_g,1Mb_^2^ equal to its lower bound constraint of zero in disease phenotypes (solid) and simulated phenotypes with varying degrees of polygenicity and with *h*_g_^2^ matching the *h*_g–cc_^2^ of each disease (dashed). Simulation data plotted are means over 5 simulations; error bars, 95% prediction intervals assuming Bernoulli sampling variance and taking into account s.e.m. (**c**) Conservative 95% confidence intervals for the cumulative fraction of SNP-heritability explained by the 1Mb regions that contain the most SNP-heritability. Lower bounds are from a cross-validation procedure involving only the disease phenotypes while upper bounds are inferred from the empirical sampling variance of *h*_g,1Mb_^2^ estimates (Online Methods).

To understand the effect of different levels of polygenicity on the distribution of per-megabase SNP-heritability estimates, we simulated quantitative traits of varying polygenicity (2K–600K causal SNPs) with *h*_g_^2^ matching the genome-wide observed-scale *h*_g–cc_^2^ estimates for schizophrenia, dyslipidemia, and hypertension (Supplementary Table 3) using PGC2 and GERA genotypes. We then applied the same procedures we applied to the real phenotypes to obtain per-megabase SNP-heritability estimates for the simulated traits (Online Methods) and compared the simulated distributions of per-megabase estimates to the observed distributions, focusing on the fraction of 1Mb regions with *h*_g,1Mb_^2^ estimates of zero (Figure 2b). Intuitively, more polygenic traits have heritability spread more uniformly across 1Mb regions and hence have fewer *h*_g,1Mb_^2^ estimates of 0, as our simulations confirmed. (Based on this statistic, our analyses suggest that schizophrenia has a genetic architectures involving *>*20,000 causal SNPs; however, we caution that—unlike our analyses below—this estimate is contingent on our parameterization of simulated genetic architectures, as are previous estimates [11,31].)

We further interrogated our real and simulated distributions of per-megabase SNP-heritability estimates to obtain nonparametric bounds on the cumulative fraction of *h*_g_^2^ explained by varying numbers of *true* top 1Mb regions—i.e., those that harbor the most SNP-heritability *in the population*—for schizophrenia, dyslipidemia, and hypertension (Figure 2c). We observed that the probability of observing an *h*_g,1Mb_^2^ estimate of zero for a given 1Mb region is a convex function of the true SNP-heritability of that region (Supplementary Figures 3 and 4), and we harnessed this observation to obtain upper bounds on the cumulative heritability explained by true top regions. To obtain lower bounds on this quantity, we applied a cross-validation procedure (similar to ref. [32]) in which we selected top regions using subsets of the data and estimated heritability explained using left-out test samples (see Online Methods). Combining the upper and lower bounds allowed us to obtain conservative 95% confidence intervals for heritability explained by top regions (Figure 2c), as we verified in simulations (Supplementary Fig. 5). In particular, we inferred that schizophrenia has an extremely polygenic architecture, with most 1Mb regions (conservative 95% CI: 71%-100%) containing nonzero contributions to the overall SNP-heritability and very little concentration of SNP-heritability into top 1Mb regions, in contrast to dyslipidemia (Figure 2c). Importantly, these bounds are not contingent on any particular parametric model of genetic architecture (Supplementary Fig. 6): this inference uses simulation data only to interrogate the sampling variance of *h*_g,1Mb_^2^ estimates, which is largely independent of the distribution of heritability across SNPs in a region (Supplementary Fig. 4) [28]. (We note that we report only conservative 95% confidence intervals—without parameter estimates—because obtaining point estimates would require assuming a parameterization of genetic architecture.) We repeated all of these analyses using 0.5Mb regions and observed no qualitative differences in the results (Supplementary Figures 2, 3, and 7 and Supplementary Table 7).

Having computed per-megabase *h*_g,1Mb_^2^ estimates, we checked for correlations between estimated *h*_g,1Mb_^2^ and genomic annotations that vary slowly across the genome. Specifically, we tabulated GC content, genic content [6], replication timing [33], recombination rate [34], background selection [35], and methylation QTLs [36] per megabase of the genome. (Each of these annotations had an autocorrelation across consecutive 1Mb segments of at least 0.3; see Supplementary Table 9.) For each of schizophrenia, dyslipidemia, and hypertension, we observed the greatest correlation with GC content (*p <* 10^−5^) (Supplementary Table 10). We also observed significant correlations of per-megabase *h*_g,1Mb_^2^ with genic content, replication timing and recombination rate; however, upon including GC content—which is correlated with each of the other annotations (Supplementary Table 11)—as a covariate, all other correlations became non-significant (Supplementary Table 10). To further investigate this finding, we stratified 1Mb regions into GC content quintiles and partitioned SNP-heritability across these strata, observing a clear enrichment of heritability with increasing GC content (Figure 3), which we verified was not due to systematic differences in SNP counts or MAF distributions across GC quintiles (Supplementary Table 12 and Supplementary Fig. 8) and not explained by differences in meQTL counts (Supplementary Fig. 9). To quantify this enrichment, we performed finer partitioning into 50 GC content strata and regressed SNP-heritability estimates against GC content (Online Methods). We found that a 1% increase in GC content (relative to the median) corresponded to 1.0%, 4.4%, and 3.2% increases in heritability explained (relative to the means) for schizophrenia, dyslipidemia, and hypertension (95% confidence intervals, 0.3–1.6%, 2.1–6.7%, and 1.8–4.6%). Once again, repeating these analyses using 0.5Mb regions produced no qualitative differences in results (Supplementary Fig. 10 and Supplementary Tables 10 and 11). We also observed that including 10 principal component covariates per variance component or applying extremely stringent QC had negligible impact on our results (Supplementary Table 13). Likewise, repeating our analyses using PCGC regression instead of BOLT-REML produced consistent results with slightly larger standard errors (Supplementary Table 13).

**Figure 3.**
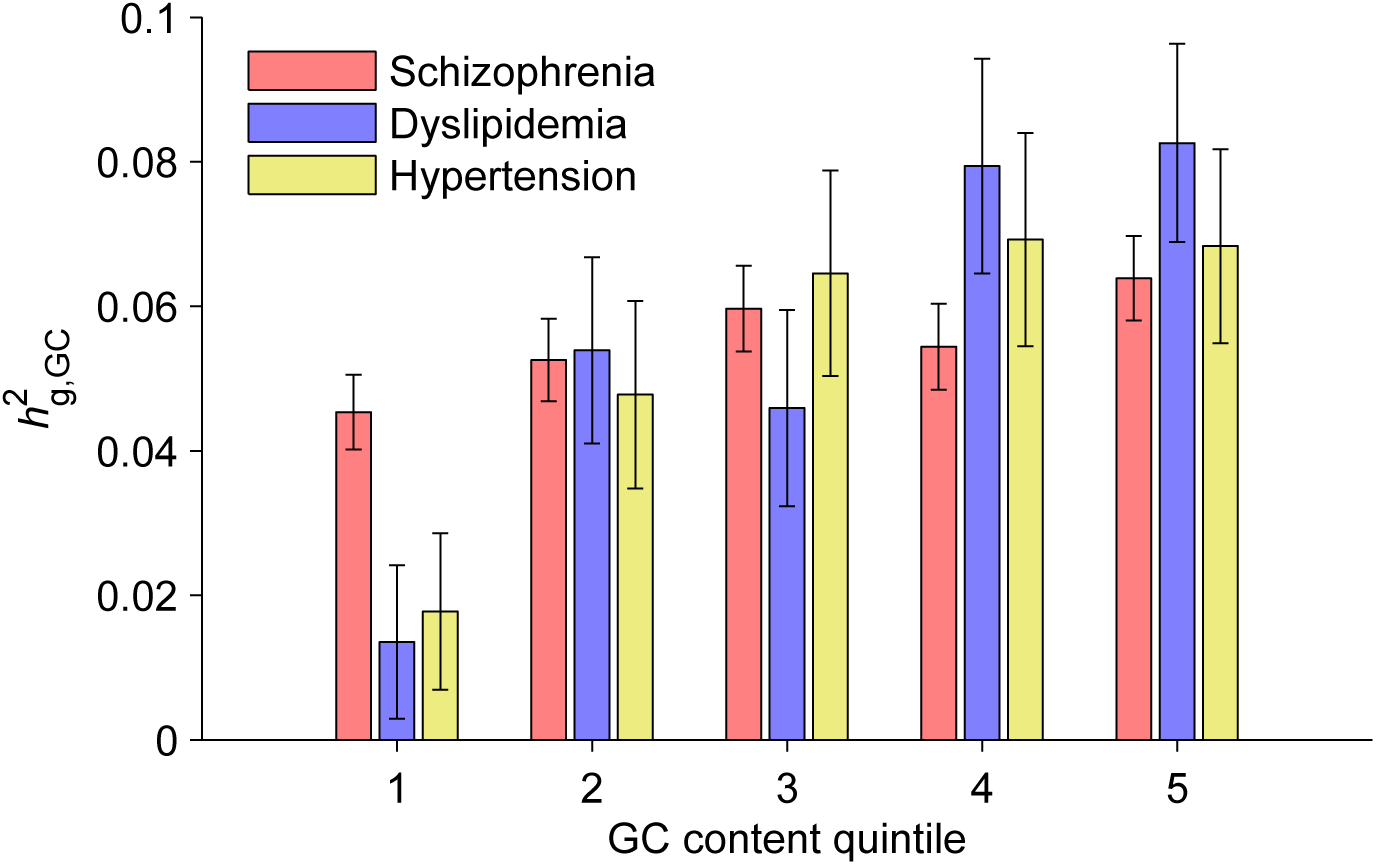
SNP-heritability of disease liabilities partitioned by GC content. GC content was computed at 1Mb resolution, after which 1Mb regions were stratified into GC quintiles for variance components analysis. Quintiles 1–5 have median GC contents of 35.7%, 38.1%, 40.2%, 42.8%, and 47.2%, respectively. Error bars, 95% confidence intervals based on REML analytic standard errors.

Finally, we performed chromosome partitioning of SNP-heritability for each disease, as previously done for schizophrenia using *N* =21K samples [5]. We confirmed a strikingly linear relationship between SNP-heritability of schizophrenia explained per chromosome and chromosome length (Supplementary Fig. 11), consistent with a highly polygenic disease architecture. In contrast, the trend for dyslipidemia was noticeably less linear, consistent with the existence of large-effect loci (Supplementary Fig. 11).

### Enrichment of SNP-heritability in higher-frequency SNPs

Given the high observed-scale heritability of schizophrenia on the full *N* =50K data set (Supplementary Table 3), we reasoned that analyses partitioning schizophrenia SNP-heritability by allele frequency would produce results with small enough standard errors to yield high-confidence conclusions, providing greater resolution than the results of ref. [5] based on *N* =21K samples. We began by running minor allele frequency (MAF)-partitioned heritability analyses of simulated quantitative phenotypes based on UK10K sequencing data (see Online Methods and URLs). We simulated genetic architectures in which causal SNPs were drawn from SNPs with MAF *p≥*0.1% and were randomly assigned allele effect sizes with variances proportional to (*p*(1 − *p*))^*α*^ for various values of *α* between –1 and 0 (ref. [28,29]) (Online Methods). Under this parameterization, *α* = –1 corresponds to a model in which rare SNPs have larger per-allele effects, so that all SNPs have the same expected contribution to variance [1], while *α* = 0 corresponds to a model with no selection [37] in which all alleles have similar per-allele effects, so that on average rarer SNPs contribute less variance. We performed MAF-partitioned analyses [29] over six MAF bins (partitioning the 2–50% MAF range) using tag SNPs from the PGC2 data set, and we observed that the heritability captured by tag SNPs in each bin (*h*_g,MAF_^2^, defined in Online Methods) accounted for most but not all of the true heritability contributed by causal UK10K variants in each bin (*h*_g,MAF_^2^, defined in Online Methods) (Fig. 4a).

**Figure 4.**
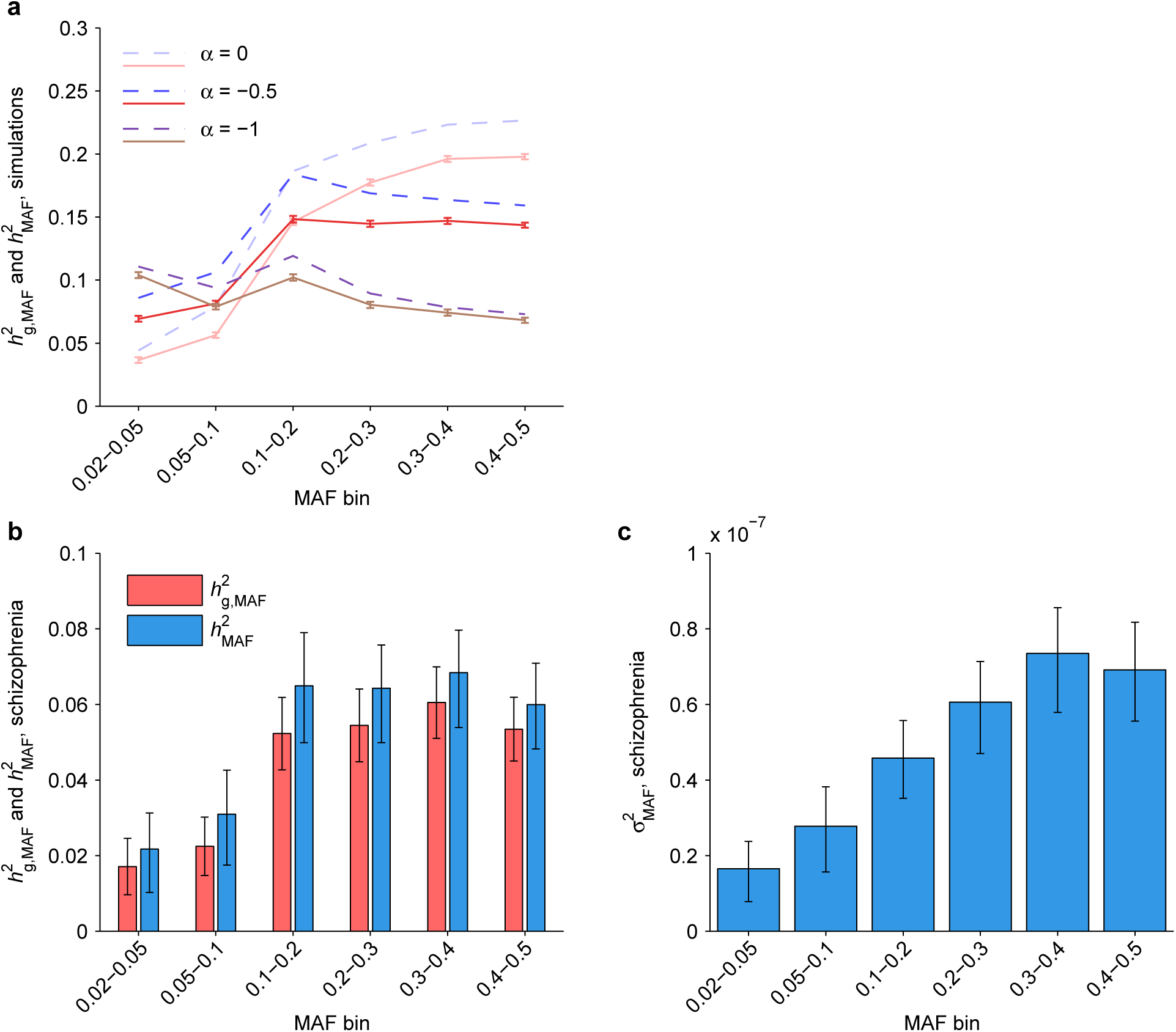
Inferred heritability of schizophrenia liability due to SNPs of various allele frequencies. (**a**) Simulated narrow-sense heritability per MAF bin (*h*_MAF_^2^, dashed blue curves) and estimated SNP-heritability per MAF bin (*h*_g,MAF_^2^, solid red curves) for quantitative phenotypes with genetic architectures in which SNPs of minor allele frequency *p* have average per-allele effect size variance proportional to (*p*(1 *− p*))^*α*^. Simulations used causal SNPs with MAF *≥* 0.1% in UK10K sequencing data and tag SNPs from our PGC2 analyses; error bars, 95% confidence intervals based on 4,000 runs. (**b**) SNP-heritability (red) and inferred narrow-sense heritability (blue) of schizophrenia liability partitioned across six MAF bins. Point estimates of narrow-sense heritability per bin are based on interpolated values of the ratio *h*_g,MAF_^2^/*h*_MAF_^2^ at *α* = –0.28, which provided the best weighted least-squares fit between observed *h*_g,MAF_^2^ and interpolated *h*_g,MAF_^2^ from the simulations in panel (a) (Supplementary Fig. 12). (**c**) Inferred narrow-sense heritability of schizophrenia liability explained per SNP in each MAF bin, i.e., *h*_MAF_^2^ in panel (b) normalized by UK10K SNP counts (Supplementary Table 14). Schizophrenia *h*_g,MAF_^2^ error bars, 95% confidence intervals based on REML analytic standard errors. Schizophrenia *h*_MAF_^2^ and *σ*_MAF_^2^ error bars, unions of 95% confidence intervals assuming –1 *≤ α ≤* 0.

We next performed MAF-partitioning of schizophrenia *h*_g_^2^ by running BOLT-REML on the full PGC2 data set with variance components corresponding to the same six MAF bins (Fig. 4b). We then estimated total narrow-sense heritability contributed per MAF bin, *h*_MAF_^2^ (Fig. 4b), by performing an inverse-variance weighted least-squares fit of observed *h*_g,MAF_^2^ against data from our simulations, interpolated for –1 *≤ α ≤* 0; this procedure yielded a best-fit value of *α* = –0.28 (jackknife s.e.=0.09) (Supplementary Fig. 12), from which we inferred *h*_MAF_^2^. To keep our inferences robust to model parameterization, we computed conservative 95% confidence intervals for *h*_MAF_^2^ (independent of the best-fit *α*, which is not our focus here) by taking the union of 95% confidence intervals assuming different values of *α* (–1 *≤ α ≤* 0). Finally, we divided *h*_MAF_^2^ by the number of UK10K SNPs per bin (Supplementary Table 14) to estimate the average heritability explained per SNP in each MAF bin, *σ*_MAF_^2^ (Fig. 4c), observing a clear increase in heritability explained per SNP with increasing allele frequency. Repeating the MAF-partitioning using PCGC regression produced consistent results with slightly larger standard errors (Supplementary Table 13). We observed the same general trend in analyses of GERA diseases, although the results were noisier due to smaller *h*_g–cc_^2^ (Supplementary Fig. 13).

### Genetic correlations across GERA diseases

The availability of multiple phenotypes across all GERA samples also allowed us to estimate the genetic correlations and total correlations (*r*_g_ and *r*_*l*_, defined in Online Methods) among disease liabilities (Figure 5 and Supplementary Table 15). We estimated genetic correlations by running bivariate BOLT-REML for each pair of case-control traits [7] and total liability-scale correlations by Monte Carlo simulations to match total observed-scale correlations (Online Methods). We first ran the analysis using only our standard set of covariates (age, sex, 10 principal components, and Affymetrix kit type) (Fig. 5a) and then reran the analysis including BMI as an additional covariate (Fig. 5b). We verified that of the nine survey-derived covariates provided with the GERA data set, BMI was the only one relevant to our analysis (Supplementary Fig. 14). Interestingly, we observed that adjusting for BMI lowered genetic correlations by a multiplicative factor of 0.75 (s.e.=0.05) and total correlations by a factor of 0.81 (s.e.=0.03), as assessed by regressing BMI-adjusted correlations on unadjusted correlations, suggesting that some correlation signal among these diseases may be mediated by BMI. Of the 13 significant genetic correlations in the unadjusted analysis, six became non-significant upon adjusting for BMI, leaving a very strong genetic correlation between asthma and allergic rhinitis (*r*_g_=0.85, s.e.=0.11) and a cluster of six moderate genetic correlations among cardiovascular disease, type 2 diabetes, dyslipidemia, and hypertension (*r*_g_=0.27–0.43) (Supplementary Table 15).

**Figure 5.**
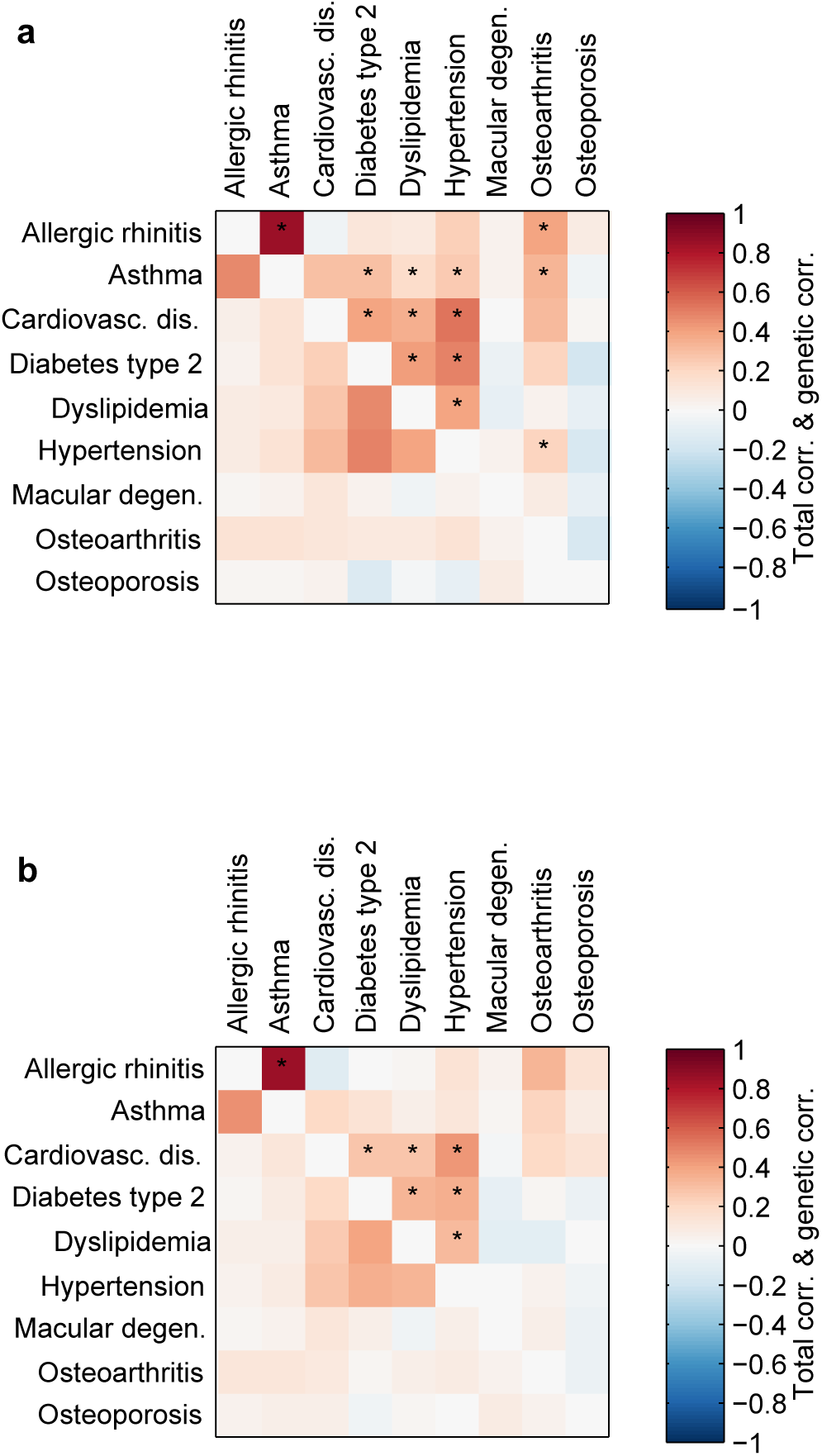
Genetic correlations and total correlations of GERA disease liabilities. (**a**) Correlations from bivariate analyses using only age, sex, 10 principal components, and Affymetrix kit type as covariates. (**b**) Correlations from bivariate analyses including BMI as an additional covariate. Genetic correlations are above the diagonals; total liability correlations are below the diagonals. Asterisks indicate genetic correlations that are significantly positive (*z >* 3) accounting for 36 trait pairs tested. Numeric data including standard errors are provided in Supplementary Table 15.

We further investigated the relationship between genetic correlations (*r*_g_) and total correlations (*r*_*l*_) among disease liabilities. We observed that *r*_g_ significantly exceeded *r*_*l*_ for asthma and allergic rhinitis (*r*_g_=0.85 vs. *r*_*l*_=0.46; *p*=0.008) after adjusting for 36 hypotheses tested; no other pair reached significance. We also observed an approximately linear relationship between genetic correlation and total liability correlation; regressing *r*_g_ on *r*_*l*_ yielded a proportionality constant of *r*_g_/*r*_*l*_=1.3 (s.e.=0.1, with the caveat that the 36 trait pairs are not independent) robust to the choice of whether or not to use BMI as a covariate (Supplementary Fig. 15).

## Discussion

We have introduced a new fast algorithm, BOLT-REML, for variance components analysis involving multiple variance components and multiple traits, and demonstrated that it enables accurate large-sample heritability analyses that were previously computationally intractable. Such analyses will be essential to attaining the statistical resolution necessary to reveal deeper insights into the genetic architecture of complex traits (Supplementary Table 16) [15,16]. We have applied BOLT-REML to study human complex diseases in roughly 50K samples from each of the PGC2 and GERA data sets. At this sample size, we uncovered multiple insights into complex disease architecture, including extreme polygenicity of schizophrenia, enrichment of complex disease SNP-heritability in GC-rich regions and in higher-frequency SNPs, and significant genetic correlations among several GERA diseases.

Our per-megabase analyses of SNP-heritability in schizophrenia, dyslipidemia, and hypertension revealed contrasting levels of polygenicity, with schizophrenia exhibiting an exceptionally polygenic architecture. Our inference that the large majority of 1Mb regions of the genome (71– 100%) contain schizophrenia loci evokes the concern that complex-trait GWAS of increasing sample sizes will ultimately implicate the entire genome, becoming uninformative [38]. Recent very large-scale GWAS [12,32,39] have begun grappling with this problem by focusing on biological pathways or gene sets instead of individual SNPs [40]. While previous studies have provided evidence for a highly polygenic architecture for schizophrenia [9,41], no previous study has provided a quantification of the extreme level of polygenicity we have observed here; in light of this result, methods that further interrogate association signal at the pathway level will be essential to extracting further biological insights about schizophrenia [42]. An additional question that this finding raises is whether the polygenicity would diminish in analyses with more homogeneous sample recruitment or phenotype (e.g., treatment resistant); future studies may be sufficiently powered to answer this question. As to our observation of enrichment of SNP-heritability with increasing GC content, further study will be required to disentangle the mechanisms underlying this phenomenon; previous work has shown that GC architecture has complex effects on recombination and replication timing [33] as well as DNA methylation [43].

Our results from partitioning the SNP-heritability of schizophrenia and GERA diseases across the 2–50% allele frequency spectrum shed light on the extent to which rarer SNPs tend to have larger per-allele effects, as predicted by evolutionary models [44,45]. Our analysis of schizophrenia, based on well-imputed SNPs with MAF*≥*2%, does not assess the contribution of rare variants (MAF*<*1%) due to the need for stringent QC in heritability analyses of ascertained case-control cohorts [3]; however, the trend for SNPs with MAF 2–50% (Fig. 4b,c) strongly suggests that rarer SNPs have larger effect sizes per allele, yet explain less variance per SNP. While further study of more phenotypes and rarer variants is needed, this observation implies that the implicit assumption of *α* = –1 made by standard analyses of heritability [1] and mixed model association [20,27] may be suboptimal, leaving room for further improvement on both fronts.

Our correlation analyses of GERA disease phenotypes identified a very strong genetic correlation (*r*_g_=0.85, s.e.=0.11) between asthma and allergic rhinitis. While the link between asthma and allergy has long been known and recent GWAS have identified many shared associations, the extent to which these two diseases are genetically related has not previously been quantified [46–48]. Among other disease pairs, our observation of significant genetic correlations among metabolic diseases confirms and adds resolution to previous estimates [49,50], while our observation of significant broad decreases in genetic and total correlations upon including BMI as a covariate highlights the importance of carefully considering the effects of heritable covariates when conducting and interpreting genetic analyses [51]. Additionally, our empirical observation of an approximately linear relationship between correlations of total liability and genetic correlations [52], viewed in conjuction with a similar (but noisier) empirical observation among a set of seven quantitative metabolic traits [50], suggests the generality of such a trend for human complex traits.

Methodologically, while the variance components (REML) approach [1] that we have applied and accelerated here enjoys widespread use, three alternative approaches to heritability analysis (with various trade-offs) have recently been proposed. First, the Bayesian sparse linear mixed model [53] adapts the variance components approach to better model traits with large-effect loci, slightly reducing standard errors at the expense of much larger computational cost; integrating this approach into BOLT-REML is a potential future direction. Second, PCGC regression [17], which generalizes Haseman-Elston regression [54], is not subject to downward bias under case-control ascertainment; we therefore recommmend PCGC regression for the purpose of estimating genomewide *h*_g_^2^ in such situations. (For partitioning SNP-heritability across subsets of SNPs, PCGC estimates have slightly higher standard errors than REML.) Third, LD Score regression [49,55] is a very different approach that makes inference using only GWAS summary statistics—not genotype data. LD Score regression has the disadvantage of somewhat higher standard errors (vs. REML) that further increase if inference is desired for small regions of the genome; as such, we are not currently aware of a method for assessing degree of polygenicity using summary statistics. All of these methods have the limitation that they assume independence of genetic and environmental effects; violation of this assumption may cause bias.

Compared to existing REML methods, the BOLT-REML algorithm we have proposed is much more computationally efficient; however, our approach does have limitations. First, because BOLT-REML achieves its speedup by avoiding direct computation of likelihoods, it is unable to compute likelihood ratio tests to assess whether variance parameters are significantly nonzero. In fact, the assumptions underlying REML analytic standard errors break down for parameter estimates of zero (and more generally, at the parameter space boundary; see Supplementary Note). GCTA [2] provides an unconstrained optimization feature that allows negative variance estimates, thereby sidestepping this issue and also reducing constraint-induced bias; incorporating such a feature into BOLT-REML is a potential future direction. Second, BOLT-REML, like all REML algorithms, occasionally fails to converge when variance parameters are poorly constrained, typically for multicomponent models at small sample sizes (*N ≪*5,000). Given that sample sizes are steadily increasing, however, we expect BOLT-REML to be a robust choice for harnessing the full power of large-scale cohorts to further elucidate complex trait architectures.

**URLs.** BOLT-REML software and source code (implemented in the BOLT-LMM v2.1 package), http://www.hsph.harvard.edu/alkes-price/software/.

GCTA software, http://www.complextraitgenomics.com/software/gcta/.

PCGC regression efficient software, http://github.com/gauravbhatia1/PCGCRegression.

PLINK2 software, http://www.cog-genomics.org/plink2.

KING software, http://people.virginia.edu/∼wc9c/KING/.

EIGENSOFT v6.0.1, including open-source implementation of FastPCA, http://www.hsph.harvard.edu/alkes-price/software/.

GERA data set, http://www.ncbi.nlm.nih.gov/projects/gap/cgi-bin/study.cgi?study_id=phs000674.v1.p1.

UK10K project, http://www.uk10k.org/.

## Acknowledgments

We are grateful to K. Galinsky, T. Hayeck, P. Palamara, J. Listgarten, V. Anttila, S. Sunyaev, D. Howrigan, R. Walters, P. Sullivan, M. Keller, M. Goddard, P. Visscher, J. Yang, S. Ripke, D. Golan, and S. Rosset for helpful discussions. This research was supported by US National Institutes of Health grants R01 HG006399 and R01 MH101244 and US National Institutes of Health fellowship F32 HG007805. H. K. F. was supported by the Fannie and John Hertz Foundation. Members of the Schizophrenia Working Group of the Psychiatric Genomics Consortium are listed in the Supplementary Note. Statistical analyses of PGC2 data were carried out on the Genetic Cluster Computer (http://www.geneticcluster.org) hosted by SURFsara and financially supported by the Netherlands Scientific Organization (NWO 480-05-003 PI: Posthuma) along with a supplement from the Dutch Brain Foundation and the VU University Amsterdam. Analyses of GERA data were conducted on the Orchestra High Performance Compute Cluster at Harvard Medical School, which is partially supported by grant NCRR 1S10RR028832-01.

## Online Methods

### BOLT-REML algorithm

The overall framework of the BOLT-REML algorithm is Monte Carlo AI REML [23], a Newton-type iterative optimization of the (restricted) log likelihood with respect to the variance parameters sought. BOLT-REML begins a multi-variance component analysis by computing an initial estimate of each parameter using the single variance component estimation procedure of BOLT-LMM [20] (which is the only analysis possible with BOLT-LMM). Then, in each iteration, BOLT-REML rapidly approximates the gradient of the log likelihood using pseudorandom Monte Carlo sampling [25] and the Hessian of the log likelihood using the average information matrix [26]. BOLT-REML efficiently computes both approximations using conjugate gradient iteration [18,19] with the performance optimizations applied by BOLT-LMM [20]. The approximate gradient and Hessian produce a local quadratic model of the likelihood surface, which we optimize within an adaptive trust region radius—key to achieving robust convergence—to obtain a proposed step. To evaluate success of the proposed step (i.e., determine whether to accept the step, whether to change the trust region radius, and whether the optimization has converged) we introduce a gradient-based approximation to the change in log likelihood achieved by the step. These procedures allow BOLT-REML to consistently achieve convergence in *≈O*(*M N*^1.5^) time; in contrast, existing multi-component REML algorithms either are less robust or require *O*(*M N*^2^ + *N*^3^) time (e.g., GCTA [2]). Details are described in the Supplementary Note.

### Accuracy of BOLT-REML variance components analysis

We verified the accuracy of BOLT-REML analysis by simulating quantitative traits with infinitesimal architectures using genotypes from subsets of the GERA data set and partitioning heritability by chromosome. On a first set of 50,000 simulations using genotypes from *N* =2,000 samples on chromosomes 21–22, BOLT-REML correctly estimated components of heritability, computing nearly identical results to GCTA [2] when run with 100 Monte Carlo trials, and incurring only 1.03 times higher standard errors when run with 15 Monte Carlo trials (Supplementary Table 17), consistent with theory (Supplementary Note). On additional sets of 100 simulations using genotypes from *N* =10,000 samples on chromosomes 1–2, BOLT-REML correctly estimated genetic correlations in bivariate analyses of simulated quantitative traits [7] (Supplementary Table 18) and randomly ascertained case-control traits using a liability threshold model [3] (Supplementary Table 19). Finally, in simulated *N* =50K case-control cohorts over-ascertained for cases (including population stratification and varying polygenicity), we observed that while absolute estimates of heritability were downward biased, as previously demonstrated [17,27], relative contributions of variance components and their standard errors were still accurately estimated when partitioning heritability by chromosome or minor allele frequency (Supplementary Figures 16–19).

### PGC2 data set

We analyzed the PGC2 schizophrenia data set [12], applying the following filters. Of 39 European-ancestry cohorts available to us for analysis, we first eliminated 10 cohorts (containing 12% of the available samples) with the lowest numbers of well-imputed SNPs. We further filtered out samples with *<*90% European ancestry as determined by SNPweights v2.0 (ref. [56]). Finally, we extracted an unrelated subset of individuals (pairwise genetic similarity *<*0.0884) using KING v1.4 `--unrelated --degree 3`; see URLs (ref. [57,58]), comprising 22,177 cases and 27,629 controls (Supplementary Table 2). Of the imputed genotypes previously computed for each cohort, we restricted to well-imputed autosomal markers (genotype call confidence *P >*0.8 with *<*2% missing rate in the cohort), given that stringent QC is critical to avoid inflated estimates of components of heritability in ascertained case-control data [3]. We then merged the 29 cohorts, taking the union of remaining markers across cohorts and then restricting to markers with total missing rate *<*5%, leaving 4.4 million markers. We further imposed a *>*2% MAF threshold based on the imputation quality of typical arrays at low MAF [59], yielding 3.9 million markers in substantial LD, to which we applied two rounds of LD-pruning at *r*^2^=0.9 (PLINK2 [60] `--indep-pairwise 50 5 0.9`; see URLs), reducing the number of markers to 596,583 and finally 472,178. Our primary motivation for pruning was to reduce susceptibility of REML *h*_g_^2^ estimation to LD bias [28–30]; additionally, pruning reduced computational costs.

### GERA data set

We analyzed GERA samples (see URLs; dbGaP study accession phs000674.v1.p1) typed on the GERA EUR chip [59] with phenotypes available for each of 22 disease conditions based on electronic medical records. (Our primary analyses did not include survey-derived phenotypes such as BMI, as the data use conditions stipulated that these phenotypes could only be used as covariates.) We applied similar filters as above, eliminating samples with *<*90% European ancestry and samples with missing sex, and extracting an unrelated subset of 54,734 individuals using PLINK2 (`--rel-cutoff 0.05`). We removed SNPs deviating from Hardy-Weinberg equilibrium (*p<*10^−6^) and SNPs with missing rate *>*2%, leaving 597,736 autosomal SNPs.

### UK10K data set

Our simulations used UK10K genotypes from sequencing data (see URLs); we merged the ALSPAC and TWINSUK cohorts, intersected marker sets and eliminated multi-allelic variants (leaving 18 million variants), and extracted 3,567 unrelated individuals using PLINK2.

### Definitions of heritability parameters

We define *h*_g_^2^ as the proportion of population variance in disease liability (assuming a liability threshold model [61]) explained by the best linear predictor using typed variants [6]. We call this quantity “SNP-heritability” [1] (although the set of well-imputed variants in our PGC2 data set included a small fraction of biallelic indels). We define *h*_g,MAF_^2^ as the proportion of population variance in disease liability explained by the subset of variants in a particular MAF range within the same best linear predictor (jointly fit using all typed variants) and define *h*_g,1Mb_^2^ and *h*_g,chr_^2^ analogously [6]. We define *h*^2^ as the total narrow-sense heritability—i.e., the proportion of population variance explained by the best linear predictor using all variants (including untyped variants)—and we define *h*_MAF_^2^ as the proportion of population variance explained by all variants in the MAF range (within a predictor using all variants). Finally, we note that we abuse notation slightly by using the above symbols to refer to both true population parameter values and estimates thereof.

### Estimating SNP-heritability of disease liabilities

We estimated *h*_g_^2^ for each GERA disease by running BOLT-REML on all samples and all markers in our filtered data set. In all our GERA analyses, we adjusted for age, sex, Affymetrix kit type, and 10 principal component (PC) covariates by residualizing genotypes and phenotypes accordingly. We included PC covariates (computed using FastPCA [62]; see URLs) to eliminate phenotypic variance explained by ancestry. We transformed raw REML parameter estimates (denoted *h*_g–cc_^2^) to *h*_g_^2^ using the linear transformation of ref. [3] assuming case fraction for each GERA disease matched population risk.

For the PGC2 data set, which is over-ascertained for schizophrenia cases, we estimated *h*_g_^2^ using PCGC regression [17] (see below) in order to avoid ascertainment-induced REML bias [17,27]. In all our PGC2 analyses, we included sex, 29 study indicators, and 10 principal components as covariates and assumed schizophrenia population risk of 1% (ref. [5,11,12]).

### Computationally efficient implementation of PCGC regression

In order to run PCGC regression on *N* =50K samples, we developed a new, efficient software implementation of PCGC regression (see URLs). The new software (i) eliminates in-memory storage of *N × N* matrices by accumulating dot products among regressors on-the-fly (i.e., streaming the genetic relationship matrix inputs); (ii) speeds up jackknife computations (by streaming the GRMs in one pass); (iii) eliminates storage of “cleaned” GRMs (i.e., GRMs with PCs projected out) by projecting PCs on-the-fly.

### Partitioning SNP-heritability across genomic regions

We estimated per-chromosome *h*_g,chr_^2^ by running BOLT-REML on all samples and markers using one variance component per chromosome and rescaling raw REML parameter estimates and standard errors by *h*_g_^2^/*h*_g–cc_^2^ (Supplementary Table 3), noting that relative variance contributions are accurately estimated by REML even under case-control ascertainment (Supplementary Figures 16–19). Estimating per-megabase *h*_g,1Mb_^2^ in an analogous manner would have required fitting a *>*2500-variance component model, which was computationally intractable, so we instead performed the computation on contiguous chromosomal segments of up to 100 regions at a time, parallelizing computations using GNU `parallel` [63]. We used joint multi-VC analyses rather than fixed effect analyses of one region at a time to improve robustness against potential confounding (e.g., subtle structure or LD between SNPs in nearby windows): any such confounding would contribute to multiple one-region-at-a-time fixed effect analyses, whereas it is spread across a joint random-effects analysis. For schizophrenia, we used one variance component per 1Mb region in the segment (discarding regions containing *<*5 markers) plus a single additional variance component containing all remaining markers. (This approach is similar to ref. [64] but computationally cheaper than directly applying ref. [64] using BOLT-REML.) Including all markers in the model was necessary because of ascertainment-induced genome-wide “linkage disequilibrium” among causal variants [27]; we observed that analyses without the all-remaining-markers variance component produced inflated estimates. For the GERA diseases, we did not observe this phenomenon, as expected for a randomly ascertained trait, so for computational efficiency we included only markers in flanking 1Mb regions in the additional variance component. We ran BOLT-REML with 15 Monte Carlo trials for the extensive computations in this section; we used 100 Monte Carlo trials in all other analyses. We note that we were unable to perform these analyses using PCGC regression due to the disk space requirements of storing 100 different 50K × 50K GRMs.

We estimated per-GC quintile *h*_g,GC_^2^ by stratifying 1Mb regions into GC quintiles and running BOLT-REML as above with one variance component per quintile. To obtain finer resolution for regression analyses, we further stratified 1Mb regions into 50 GC content strata. We then performed a series of BOLT-REML analyses with one variance component containing the first *n* strata and a second variance component containing the last 50 – *n* strata, and we estimated *h*_g,GC_^2^ of the *n*^th^ stratum as the difference between the SNP-heritability estimates for *n* and *n* – 1 strata.

### Bounding SNP-heritability explained by top 1Mb regions

We bounded the population variance in disease liability explained by the 1Mb regions with largest true *h*_g,1Mb_^2^ using the following procedure. We inferred an upper bound by analyzing the observed distribution of *h*_g,1Mb_^2^ estimates and accounting for sampling variance. Explicitly, we analyzed the probability of obtaining a zero *h*_g,1Mb_^2^ estimate, *P* (0), as a function of the actual value of *h*_g,1Mb_^2^ (relative to its mean). Because of sampling noise and the nonnegativity constraint on our REML *h*_g,1Mb_^2^ estimates, *P* (0) is always positive. In lieu of an analytic formula for *P* (0) as a function of actual *h*_g,1Mb_^2^, we obtained Monte Carlo estimates of *P* (0) by simulating quantitative traits (for the samples analyzed, using their actual genotypes) with heritability equal to the *h*_g–cc_^2^ of the actual disease status (Supplementary Table 3). We distributed heritability across varying numbers of causal variants (13 values ranging from 2,000 random markers to all available markers) and assigned each normalized causal variant a normally distributed effect size, repeating each simulation five times. For each of the 65 simulated traits, we estimated *h*_g,1Mb_^2^ for each 1Mb region. Combining this data with the actual *h*_g,1Mb_^2^ per region (i.e., the sum of squared simulated effect sizes), and aggregating the data from all simulations and all 1Mb regions, we obtained a clean empirical estimate of *P* (0) as a function of actual *h*_g,1Mb_^2^, which we observed was well-fit by a sum of two exponentials (Supplementary Fig. 3). While the empirical curve was based on simulation data, it is robust to the genetic architecture used in simulations (e.g., varying numbers of causal SNPs and normal vs. Laplace effect size distributions, Supplementary Fig. 4), as it simply measures the sampling distribution of constrained REML estimates for our genotype data at a given actual *h*_g,1Mb_^2^.

To interpret the observed fraction of zero *h*_g,1Mb_^2^ estimates in light of this information, we harnessed the fact that the decay curve of *P* (0) vs. actual *h*_g,1Mb_^2^ is convex (Supplementary Fig. 3). In particular, if a set of 1Mb regions has a fixed average actual *h*_g,1Mb_^2^, their average *P* (0) is minimized when all the regions have equal actual *h*_g,1Mb_^2^ (by Jensen’s inequality). Conversely, an uneven distribution of actual *h*_g,1Mb_^2^ across regions tends to increase the number of zero *h*_g,1Mb_^2^ estimates. These observations allowed us to bound the maximum fraction of *h*_g_^2^ that could be explained by top 1Mb regions and still be consistent with the observed fraction of zero *h*_g,1Mb_^2^ estimates. Explicitly, if a certain number of top regions explain SNP-heritability *h*_g,top_^2^, then the sum of *P* (0) over all regions is minimized by setting *h*_g,1Mb_^2^ of each top region to (*h*_g,top_^2^/ #top regions) and *h*_g,1Mb_^2^ of each remaining region to (*h*_g_^2^ -*h*_g,top_^2^) / (#non-top regions). We therefore bounded *h*_g,top_^2^ by requiring this minimum expected number of zero *h*_g,1Mb_^2^ estimates to be at most the observed number of zero *h*_g,1Mb_^2^ estimates (plus 1.96 times its s.e. for a conservative 95% confidence bound). We checked the accuracy of this procedure using simulated case-control ascertained data sets (Supplementary Fig. 5).

We obtained lower bounds on the fraction of *h*_g_^2^ explained by top 1Mb regions by 3-fold cross-validation. For each fold in turn, we estimated *h*_g,1Mb_^2^ for each region using the remaining two folds, ranked regions accordingly, and then estimated the SNP-heritability explained by top-ranked regions using the left-out fold. We repeated this procedure three times, obtaining nine estimates per fraction of regions, and computed the mean minus 1.96 times the s.d./3 as a conservative 95% confidence lower bound on SNP-heritability explained by top regions. We estimate s.e. using s.d./3 because the variance of heritability estimates scales with the number of sample pairs (*N*^2^) for *N ≪M* [15,16]. This s.e. estimate is not theoretically precise due to the complexities of sample reuse in cross-validation [65], but a rough estimate (see Supplementary Table 4 for empirical support) suffices given that the lower bound is probably a substantial underestimate (i.e., very conservative): the finite sample size of the training folds prevents an accurate ranking of regions, especially those contributing small amounts of variance.

### Partitioning SNP-heritability across allele frequency bins

We computed per-MAF bin *h*_g,MAF_^2^ estimates in a manner analogous to *h*_g,chr_^2^ estimates. To infer per-MAF bin *h*_MAF_^2^ explained by untyped as well as typed variants, we ran simulations using UK10K sequencing data to assess the tagging efficiency of our PGC2 and GERA marker sets in various MAF ranges. Specifically, we simulated fully heritable quantitative traits in which normalized SNPs with MAF *p≥*0.1% (in the UK10K data) were selected as causal with probability 0.5% and assigned normally distributed effect sizes with variance (*p*(1 *− p*))^*α*^. (This setup assumes that UK10K SNPs explain all narrow-sense heritability, but given that we are only interested in tagging efficiency at MAF*≥*2%, our estimation procedure is robust to violations of this assumption. We also note that our choice of a normal distribution of effect sizes is inconsequential given the robustness of REML estimates to a wide range of genetic architectures [28].) We performed 4,000 simulations for each of *α* = 0, –0.25, –0.5, –1. For each marker set, we then computed REML estimates of *h*_g,MAF_^2^ for each simulated trait across six MAF bins (Fig. 4) using one variance component per bin [29] and restricting to SNPs in the marker set. A small subset of the PGC2 marker IDs (8%) and GERA SNP IDs (4%) were not present among the UK10K SNP IDs, so we did not include these markers in our REML analyses of simulated traits; we verified that the inclusion vs. exclusion of these markers had a negligible effect on schizophrenia *h*_g,MAF_^2^ estimates (Supplementary Fig. 20). We performed REML analyses of UK10K simulated traits using a slightly modified version of GCTA v1.21 [2] in order to perform robust unconstrained REML (i.e., allow negative *h*_g,MAF_^2^ estimates); at low sample sizes, constrained REML estimates are upward biased due to noise and the positivity constraint. (We modified GCTA to improve robustness in this setting by adding a trust region framework to its REML optimization.) Finally, we computed *h*_MAF_^2^ for the simulated traits by summing squared simulated effect sizes.

### Estimating genetic correlations and total correlations of disease liabilities

For each pair of GERA diseases, we estimated their genetic correlation (denoted *r*_g_) directly from bivariate BOLT-REML, which models both genetic and residual covariance, using all samples and markers. Under a liability threshold model, the estimated genetic correlation (using observed case-control phenotypes) accurately reflects the genetic correlation of underlying disease liabilities, so we did not need to transform raw BOLT-REML *r*_g_ parameter estimates [7]. However, the total correlation of observed case-control phenotypes is damped relative to the total correlation of underlying disease liabilities (which we denote by *r*_*l*_): assuming two diseases have bivariate normal liabilities *l*_1_ and *l*_2_ with correlation *r*_*l*_, the correlation of case-control phenotypes is *r*_p_= corr(*l*_1_*>z*_1_, *l*_2_*>z*_2_), where *z*_1_ and *z*_2_ are appropriate liability thresholds. In general, *|r*_p_*|≤|r_l_|* under a bivariate normal liability threshold model; e.g., two traits with the same liabilities (*r*_*l*_=1) but different thresholds (*z*_1_≠*z*_2_) have *r*_p_*<r_l_*. We recovered *r*_*l*_ from *r*_p_ by straightforward Monte Carlo simulation, performing a binary search to determine the value of *r*_*l*_ producing the observed *r*_p_ assuming values of *z*_1_ and *z*_2_ corresponding to GERA case fractions. Similarly, we obtained an s.e. for *r*_*l*_ by transforming the 95% confidence interval for *r*_p_ (based on its s.e. of 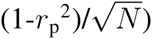 in the same way. Finally, we note that for analyses in which we included BMI (coded on a 1–5 scale in the GERA data) as a covariate, we included an additional missing indicator covariate marking samples with missing BMI (5%).

